## Supplementary Information for "Automated systematic evaluation of cryo-EM specimens with SmartScope"

### SmartScope: Framework for unsupervised cryo-EM imaging

**Supplementary Information**

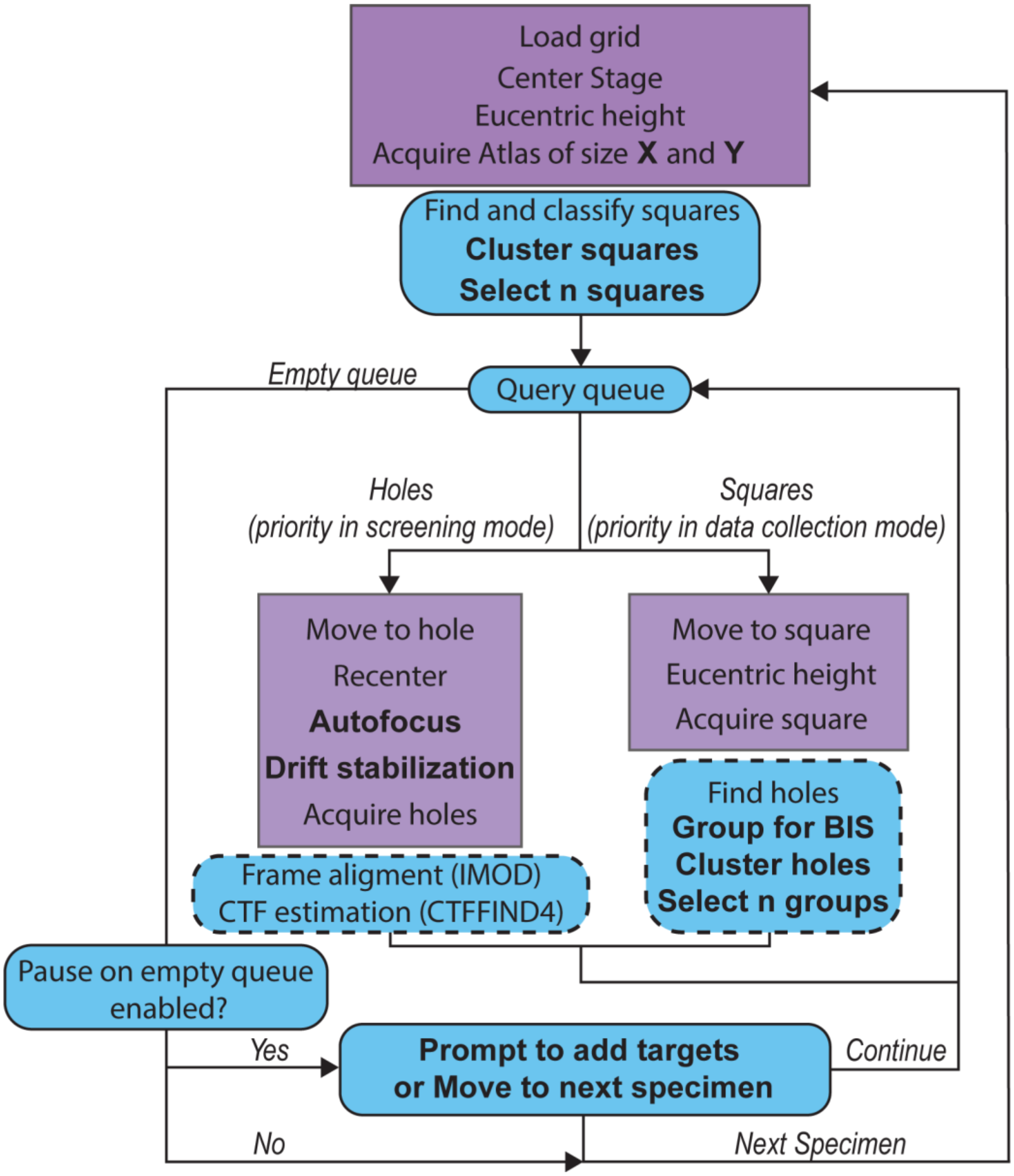


**Supplementary Figure 1. Detailed SmartScope workflow.** Steps carried out by SerialEM (Mastronarde, 2005) are shown in purple and steps carried out by SmartScope are shown in blue. Steps that can be modified within the sessions setup or during the session are shown in bold (See Table S1 for descriptions of the parameters). The steps in the dashed boxes are executed asynchronously from the main workflow, meaning that the workflow moves to the next steps while these are executing.

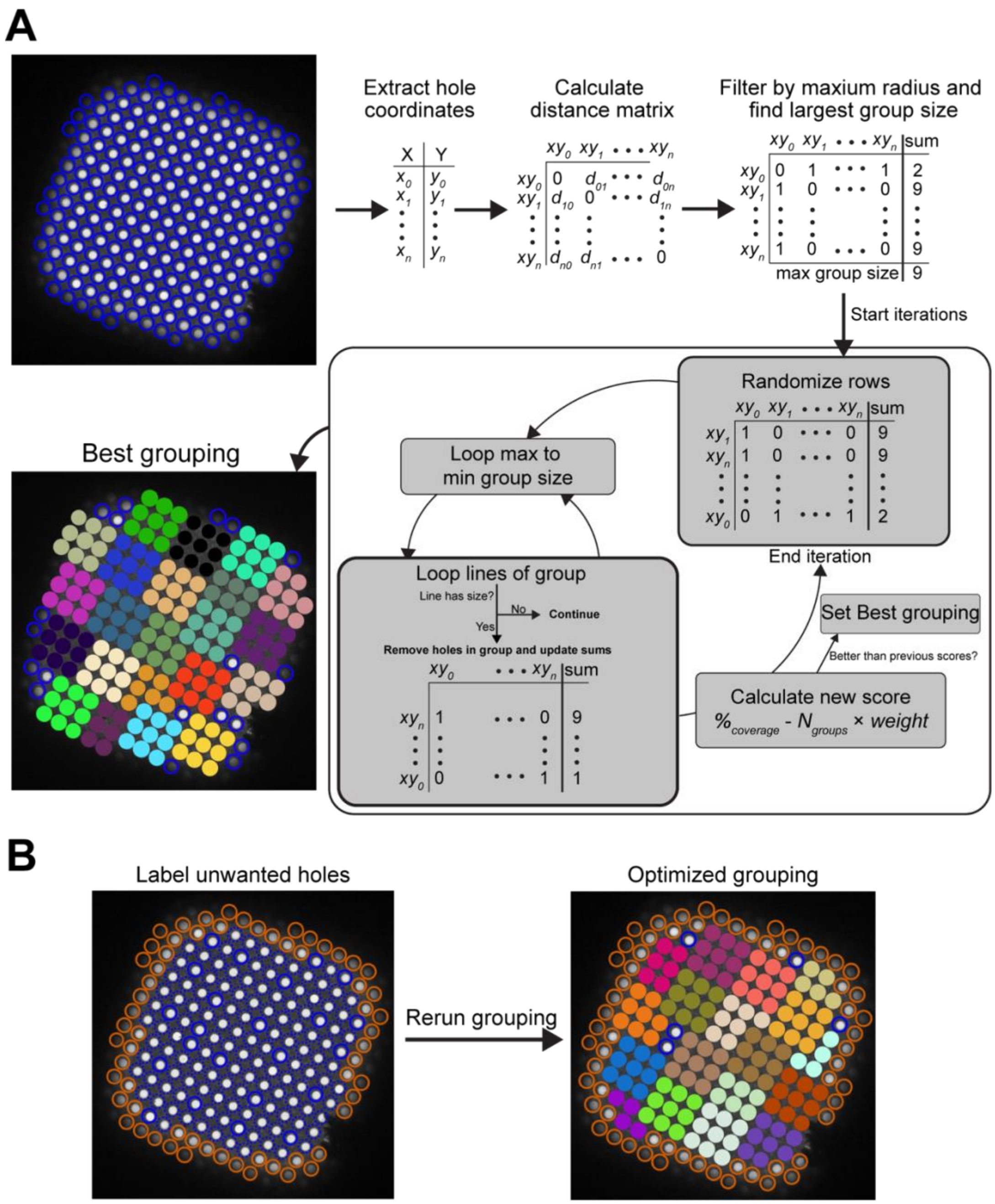


**Supplementary Figure 2. Beam-image shift hole grouping algorithm**. **A.** The hole grouping function takes the coordinates of the holes in a square, the maximum grouping radius in microns and a minimum group size. The algorithm uses the distance matrix between the holes and attempts to group them to maximize the coverage while minimizing the number of groups. Each iteration, shown in the dark gray box, uses a randomized matrix. The iterations will stop after 500, unless 250 consecutive iterations did not improve the score. Scores can be weighted in favor of coverage or number of groups by changing the weight (default weight = 2). **B** Example of how grouping can be optimized by labeling holes bad holes (red) and re-running the grouping algorithm.

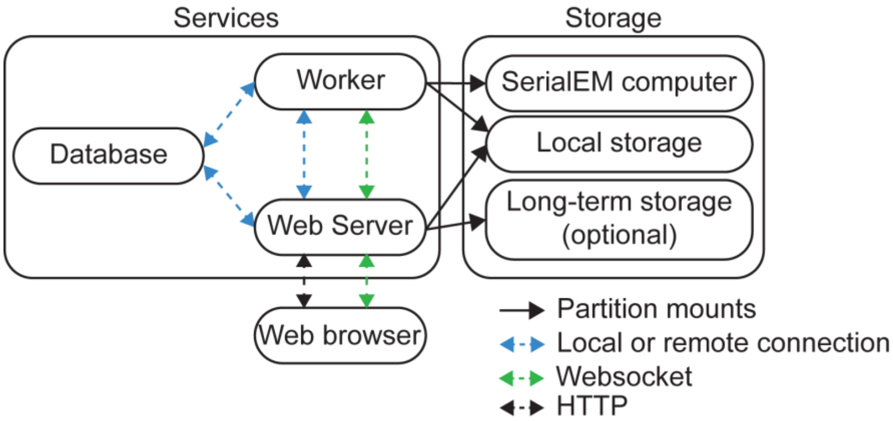


**Supplementary Figure 3. Overall software architecture of SmartScope**. SmartScope uses a database server (MariaDB), a web application with a web server and a Worker that runs the main processes. Each component can be installed on different computers and communicate with each other using Socket or SSH connections. Ongoing sessions are updated in real-time via a WebSocket connection. The worker is the only service that requires network access to the SerialEM computer. The worker and Web server need access to a shared disk to continuously update the results. After data is acquired, it can be moved to an optional long-term storage such as a local network drive or to an AWS S3 bucket.

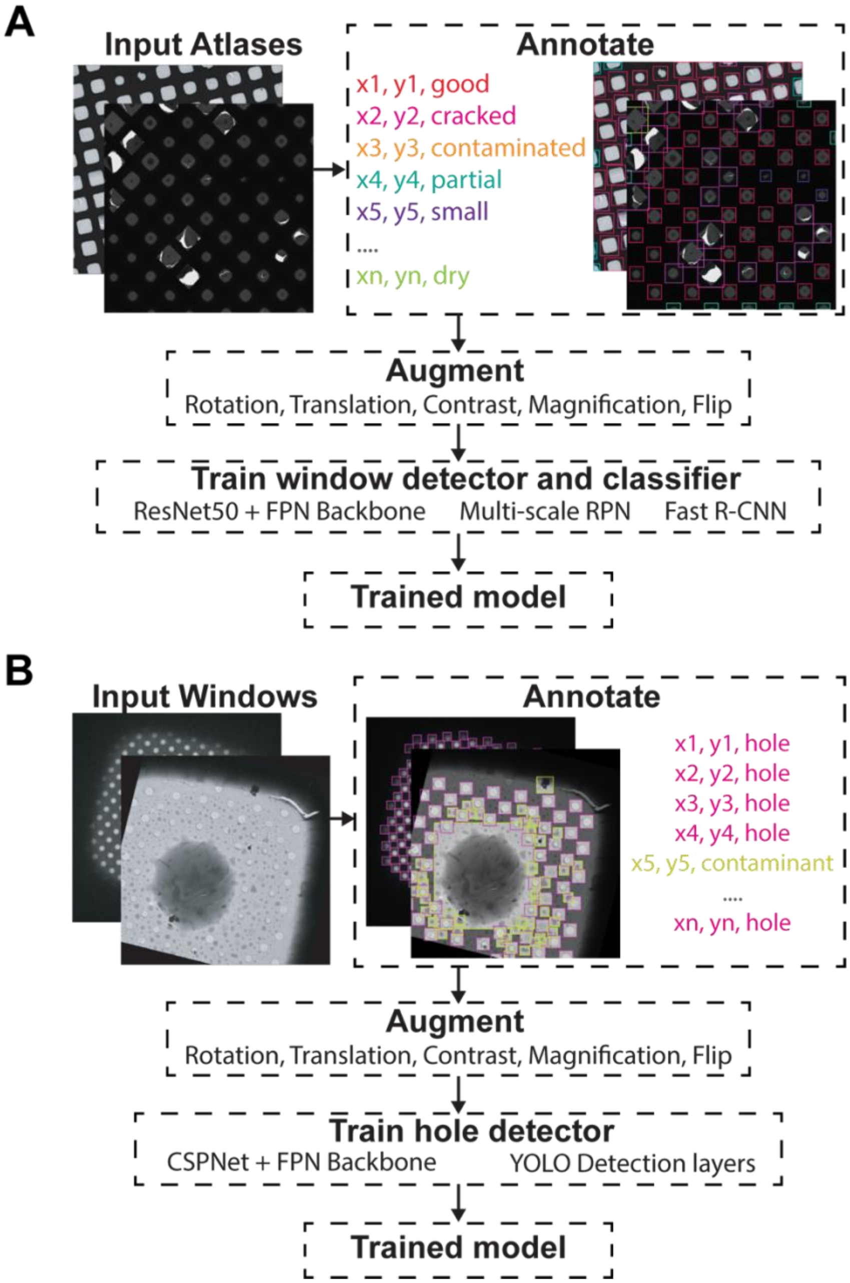


**Supplementary Figure 4. Object detection training strategy.** Atlas (**A**) and Windows (**B**) where manually picked and annotated. The annotated dataset was augmented using a combination of rotations, translations, contrast variations, magnifications and flip transforms. The models were trained against their specific architecture.

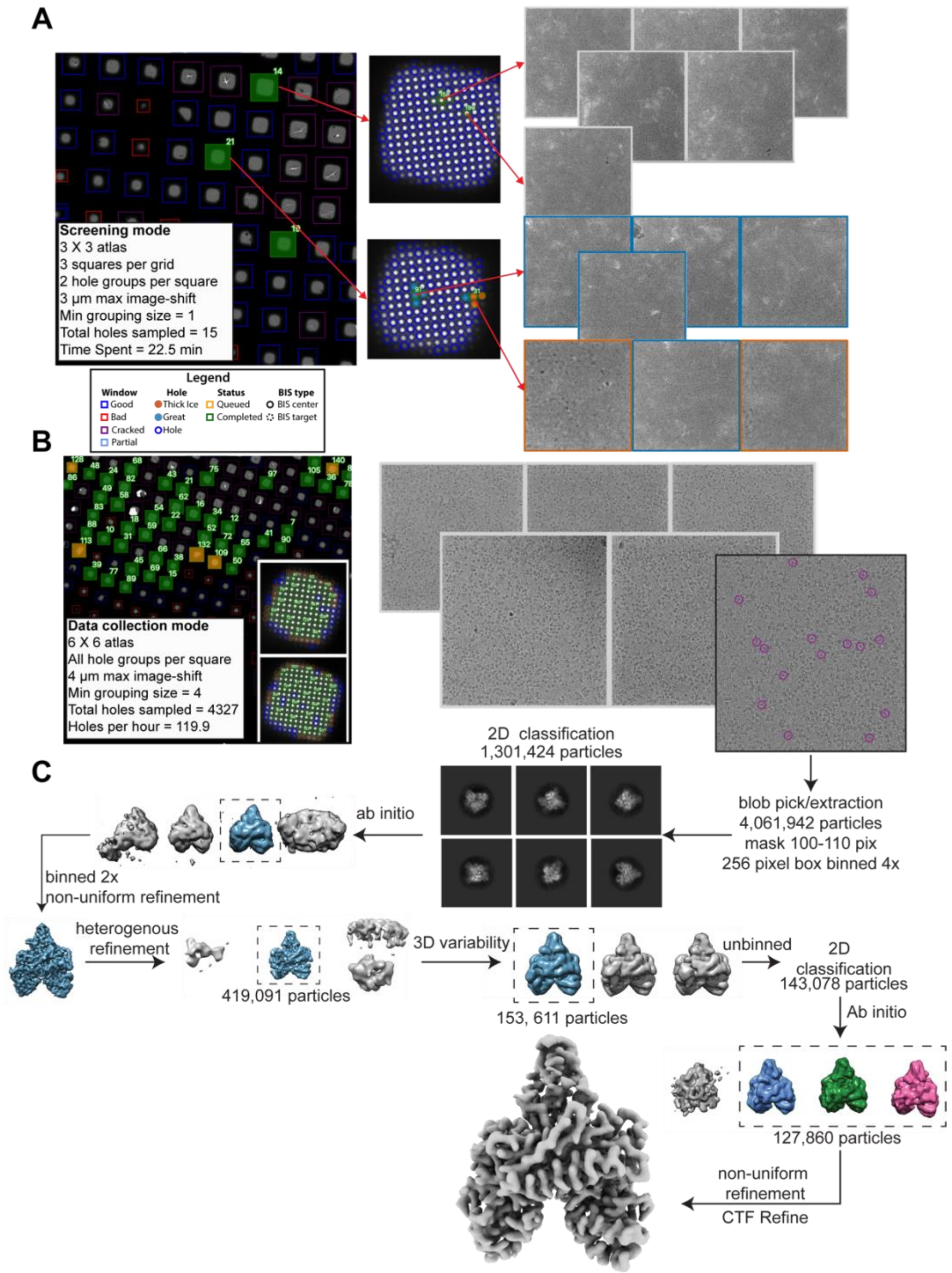


**Supplementary Figure 5. Structure determination of POLG2 using SmartScope**. **A** Screening of the specimen used for POLG2 data collection. Sample images from screening are shown. Data can be labelled from the webUI as a way to keep track of the quality. The holes will be colored according to their quality (red: Thick ice, blue: great). **B** Data collection of the same POLG2 specimen with sample squares and micrographs. Purple circles on the micrographs represent POLG2 particles (right). **C** Data processing of the POLG2 dataset using cryoSPARC (Punjani et al., 2017).

**Supplementary Table 1.** Input parameters for a SmartScope session. Parameter names and their description. This is presented as a form on the web interface. These parameters can also be updated during the imaging process.

| Parameter name | Description |
|---|---|
| General session parameters | |
| Session name | Name of the microscopy session |
| Group | Group name of the microscopy session. Usually the principal investigator's name. |
| Microscope | Microscope being used for the session |
| Detector | Detector being used for the session |
| Collection Parameters | |
| Atlas X | Number of tiles for the Atlas acquisition in X axis (default: 3) |
| Atlas Y | Number of tiles for the Atlas acquisition in Y axis (default: 3) |
| Square X | Number of tiles for the window acquisition in X axis (default: 1) |
| Square Y | Number of tiles for the window acquisition in the Y axis (default: 1) |
| Squares num | Number of windows to be selected for high-magnification imaging. (default: 3) |
| Holes per square | Number of targets per windows to be selected for higher-magnification imaging. If 0 is entered, all targets will be selected and data collection mode will be enabled. (default: 3) |
| BIS max distance | Beam-Image shift grouping radius in microns. (default: 0) |
| Min BIS group size | Smallest Beam-Image shift group size to be considered. (default: 1) |
| Target defocus min | Lower end of the defocus range for rolling defocus in microns. (default: -2) |
| Target defocus max | Higher end of the defocus range for rolling defocus in microns. (default: -2) |
| Defocus step | Step by which the defocus is varied between each target group. (default: 0) |
| Drift crit | Drift threshold to be met during the drift settling procedure before proceeding with high-magnification imaging. Use -1 to disable (default: -1) |
| Tilt angle | Tilt angle to use for high-magnification imaging. Works with BIS enabled. (default: 0) |
| Save frames | Whether to save the movie frames or return aligned sum. (default: Saving enabled) |
| Zeroloss delay | Time delay in hours for zero loss peak refinement. Only useful if the microscope has an energy filter. Use -1 to deactivate. (default: -1) |
| Autoloader (1 per grid) | |
| Name | Name of the grid |
| Position | Position in the autoloader |
| Hole Type | Grid hole spacing type (*i.e.* R1.2/1.3) |
| Mesh Size | Grid mesh size and spacing (*i.e.* 300) |
| Mesh Material | Grid mesh material (*i.e.* carbon or gold) |

**Supplementary Table 2.** Average precision obtained for each type of grid square.

| Feature type | Small | Cracked | Dry | Contaminated | Good | Partial |
|---|---|---|---|---|---|---|

| | | | | | | |
|---|---|---|---|---|---|---|
| Without Augmentation | 65.7% | 71.6% | 77.9% | 46.7% | 77.7% | 45.0% |
| With Augmentation | 73.4% | 76.5% | 79.4% | 50.41% | 81.2 % | 45.5% |

**Supplementary Table 3. Smartscope screening mode statistics**. All grids were collected using a K2 detector. The default parameters for screening mode are a 9-tile atlas, 3 squares and 3 holes per square.

| Total specimens = 981 | min | max | mean | median | standard deviation |
|---|---|---|---|---|---|
| Squares sampled | 0 | 7 | 2.5 | 3.0 | 1.3 |
| Holes sampled | 0 | 33 | 8.0 | 9.0 | 5.0 |
| Time Spent on specimen (min) | 3.0 | 56.3 | 21.0 | 20.9 | 8.4 |

**Supplementary Table 4. Smartscope screening mode statistics excluding specimens with no usable or visible squares**. All grids were collected using a K2 detector. The default parameters for screening mode are a 9-tile atlas, 3 squares and 3 holes per square.

| Total specimens = 772 | min | max | mean | median | standard deviation |
|---|---|---|---|---|---|
| Square sampled | 1 | 7 | 3.0 | 3.0 | 0.9 |
| Hole sampled | 1 | 33 | 9.4 | 9.0 | 4.0 |
| Time Spent on specimen (min) | 9.1 | 56.3 | 23.0 | 21.7 | 6.7 |

**Supplementary Table 5.** Smartscope data collection mode statistics. Start times are at the start of specimen loading in the column. All grids were imaged using a K2 detector.

| Total specimens = 58 | min | max | median | mean ± SD |
|---|---|---|---|---|
| Total micrographs | 565 | 8333 | 1626 | 2155 ± 1502 |
| Holes per hour | 51 | 141 | 100 | 100 ± 22 |
| Data collection setup time (min)* | 6 | 80 | 32 | 34 ± 17 |
| Setup time per 1000 micrographs (min) | 4 | 55 | 17 | 19 ± 11 |

* Data collection times are calculated as the time needed from grid loading to the collection of the first 50 high-magnification targets minus the time required for these 50 targets to be acquired.
